# Does national power trigger ocean conservation?

**DOI:** 10.1101/2020.09.10.292045

**Authors:** Germán Baldi, Santiago A. Schauman, Patricia Gandini

## Abstract

States are reacting to the global crises of biodiversity and the provision of ecosystem services mainly through the expansion of their networks of protected areas. This reaction would have been boosted by the commitments made between the parties of the Convention on Biological Diversity, and facilitated by the opportunities offered by isolated territories, where economic interests are minimal. However, few studies have discussed the importance of national power as conservation conditionings, particularly in the ocean. In this regard, here we evaluate whether the relative extent of marine protected areas (MPAs) is related to different elements of national power. Following a quantitative approach and incorporating into analyses 155 countries, our models suggest that an increasing power (in terms of country size –land and ocean– and military capacity) is related to greater marine protection. Although these patterns could be initially associated with the ample human and economic resources of most powerful countries and with the opportunities provided by their overseas territories, different arguments would support national power elements as conservation drivers. Specifically, the exertion of such power through conservation could be linked to geopolitical strategies such as the (re)validation of a country’s sovereignty over its Exclusive Economic Zone (EEZ), the greater regulation of the circulation and use of this space, the greater influence in the regional context, and the assurance in the provision of future ecosystem goods and services. In this way, changes in geopolitical conditions could affect MPAs, compromising the effective conservation of biodiversity and ecosystem processes, as well as the sustainable management of assets.

## 1. Introduction

Oceans are notably altered due to the great pressure for their food, energy, and mineral resources [1], the presence of plastics and other pollutants [2, 3], and the effect of changes in their physical and chemical conditions [4]. Given this situation, almost all countries negotiated in 2010, within the framework of the Convention on Biological Diversity, that at least 10% of their coastal and marine territories needed to be included within networks of protected areas by 2020 [first clause of the Aichi Biodiversity Target 11, 5]. As a result, the number of marine protected areas (MPAs), as well as the protected extent, have increased significantly in the last decade, currently reaching 15,400 units and 7.3 % of global oceans [6]. This last number comes from a few extensive units of more than 100,000 km^2^ (e.g., Pitcairn Islands MPA), which occupy more than 60% of the total protected extent [7, 8]. The majority of very large MPAs are established in overseas territories, where local and corporate economic interests are minimal and where credible monitoring and conservation plans can be set [9-12]. This geographical bias in the distribution of MPAs has been attributed to the active role and interest over these areas of international Non-governmental organizations (NGOs; e.g., Conservation International, National Geographic Society, Pew Environment Group) [10, 13] and the influence of scientists and technical experts [14]. These individuals or lobby groups would set a direct line of action and communication with high-rank policymakers of countries with extensive technical and logistical capabilities [10, 14], though the relationship between national development and protection appears to be weak and far from linear [15-17].

However, the motivations underlying this sustained increase in coastal and marine conservation do not appear to be exclusively opportunistic (i.e., conservation in feasible places) nor –to a lesser extent– biocentric (i.e., the search to preserve areas of high biological diversity or sources of ecosystem services). There is a growing corpus of political/social essays that reveals the importance of geopolitical practices conditioning wildlife conservation [18] and boosting the establishment of MPAs [19-27], although these are not made explicit by conservation agencies [28]. This corpus suggests that the expansion of MPA networks helps to reaffirm or expand the state dominance over their maritime Exclusive Economic Zones (EEZ), fundamentally in core or more developed countries. Likewise, this jurisdictional progress is achieved thanks to a shift from economic and political arguments to environmental or conservationist ones.

The current state of the art of geopolitical motivations conditioning ocean conservation is based on the examination of single protected areas [e.g., 24] or single countries [e.g., 20]. Very recently, De Santo [25] broadened the political and geographical spectrums by conducting a thorough review of the long-term factors that led French, American, and British governments to the creation of very large MPAs in their overseas territories. From the sub-polar South Sandwich Islands to the sub-tropical Northwestern Hawaiian Islands, De Santo performed a survey based on scientific and grey literature that shows how most of the MPAs either harbor a rich military past or maintain a strong military presence. For the three countries, MPAs would extend their influence away from metropolises and would preserve biotic and abiotic assets for potential future consumption. Geopolitics is intimately intertwined to –or deal with– the notion of national power, and how to acquire, maintain, or increase their constitutive elements [29-31]^1^. Even though all countries generate or adopt instruments that ideally seek to strengthen their economy and increase the well-being of their population [32], or – rationally and amorally– to increase their global dominance, in the context of this paper we consider that only the most powerful ones have the capabilities of deploying geopolitical strategies in the world’s oceans [30, 33, 34].

In this sense, we quantitatively explore the effect of different elements of national power on the extent of MPAs networks. Specifically, we first evaluate whether the marine protected extent (in percentage) of each country is related to (i) the country size classification of Crowards [35, 36], (ii) the land and the ocean areas, (iii) three proxies of military capacity or power. Secondly, we evaluate whether protection is biased towards peripheral territories. We define a periphery as an EEZ geographically detached from the EEZ of the central territory, and identify if this periphery is an integral and undisputed part of a country, or is in litigation between two or more countries. We discuss the possible associations of our results with geopolitical motivations. Finally, this assessment is conducted over all countries with marine coast, therefore incorporating a wide range of economic, cultural, and biophysical conditions.

## 2. Methods

Samples of this study were the Parties of the Convention on Biological Diversity (CBD), including sovereign countries members of the United Nations, the Cook Islands and Niue (both under the Realm of New Zealand), and the United States of America (a United Nations member, but not a CBD party). We used the EEZ database from the Flanders Marine Institute [37], composed of 281 EEZ regions^2^ belonging to 156 sovereign countries^3^ (Figure 1a). We excluded from analyses 22 EEZ regions of “joint regime” or “undetermined” categories, as well as three ones belonging to the State of Palestine (a CBD Party) and Taiwan. The geographical data about marine protected areas (MPAs) was obtained from the World Database on Protected Areas, version March 2019^4^ (Figure 1b) [39]. We included all MPAs (codes “marine” = ‘1’ –”mixed areas”, or “marine” = ‘2’ –”marine areas”) categorized as I-VI under the IUCN guidelines [40], as well as “Not Reported”. We excluded all MPAs with a “Proposed” status. As we included polygons and points, we created a circular buffer around each point with the reported area. Due to a potential overestimation of the national protected extent from overlapping problems [41], all polygons were dissolved and transformed into a raster with a pixel size of 250 m. We then intersected the EEZ database with the MPA raster in order to calculate the protected extent of each EEZ unit. As there are discrepancies between the WDPA and Flanders Marine Institute databases concerning international limits, we manually discarded those sections of MPAs that exceeded the boundaries of the country to which they belong (e.g., the French Nord Bretagne MPA overlaps the Guernsey EEZ). After this data manipulation, the total global protected extent was 29.5 M km^2^, which is equivalent to 8.2 % of the oceanic surface, divided into approximately 12,400 units.

**Figure 1.**
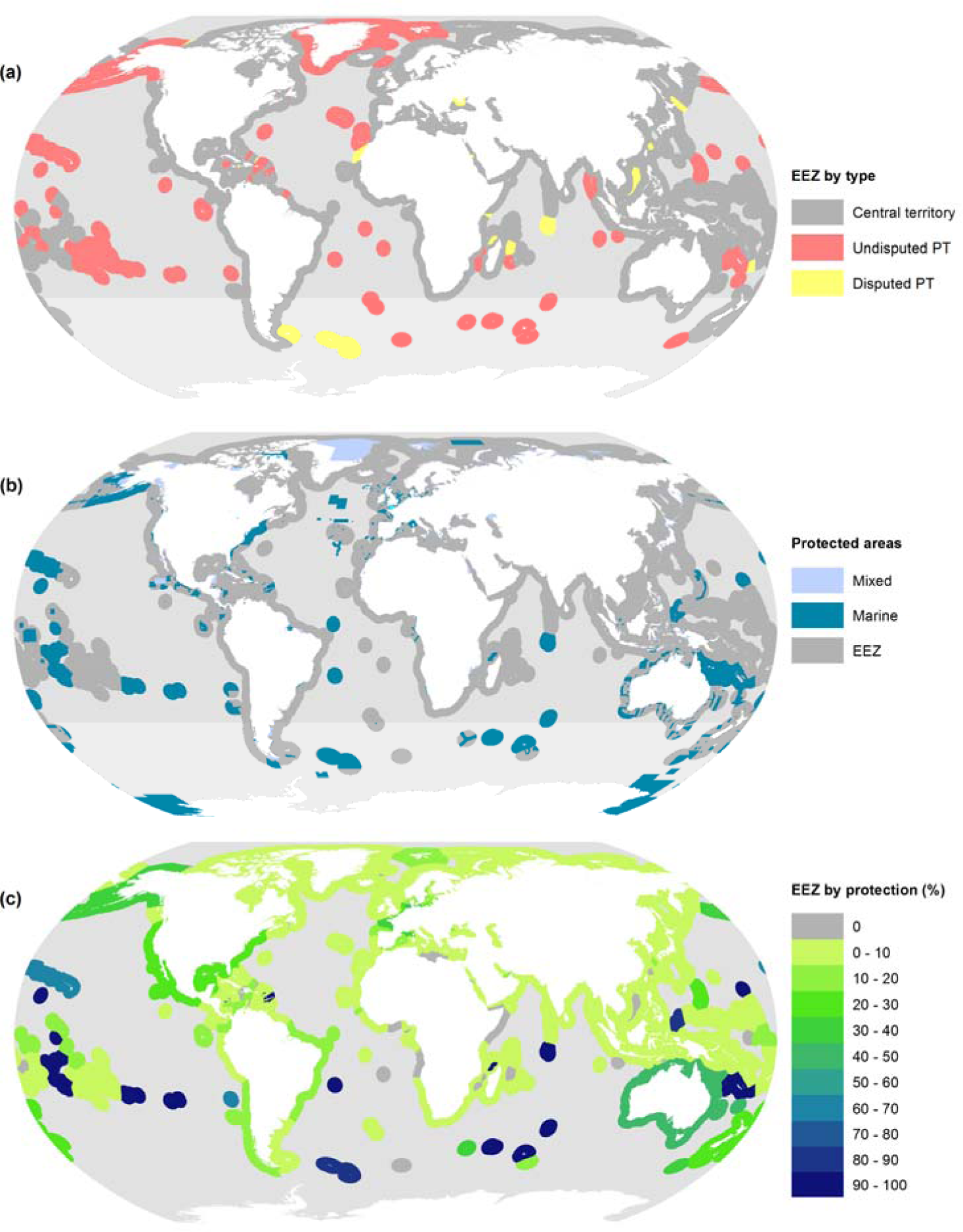
(a) Exclusive Economic Zones (EEZs) of central, disputed and undisputed peripheral territories (n = 256). (b) Protected areas categorized as I-VI and Not Reported under the International Union for Conservation of Nature guidelines [IUCN, 40]. (c) Protected extent of the EEZ according to (b).

We related the protected extent at a national level to six independent variables representing different elements of national power. The first one, Crowards’ size, is a classification of countries into five categories (from micro to very large) based on a non-hierarchical cluster analysis considering the land area, GDP, and population [35, 36]. The categorization of unclassified countries (e.g., Australia, Canada) was completed following his initial methodological scheme. The second and third variables were the land and the ocean (i.e., EEZ) areas, which depict the abundance and diversity of natural assets, influencing population and economic structure, among many other elements [29]. The next three variables represent military power, i.e., the capability to dissuade and/or coerce external entities/policies to either protect and/or advance national interests at national to global levels. Military power was depicted by the military expenditure, the Composite Indicator of National Material Capabilities (CINC) v. 5.0 [42], and the multiplicative interaction among GDP and GDP per capita [43]. Military expenditure data came from the average values 2007-2016 of the World Bank [44]. Missing cases (e.g., Comoros), came from GlobalSecurity.org [45]. The Composite Index of National Capability (CINC) is one of the most accepted method for measuring national hard power, combining data on military spending, troops, population, urban population, iron and steel production, and energy consumption. According to Beckley [43], the military expenditure and the CINC would exaggerate the wealth and military capabilities of poor, populous countries, and therefore proposed the GDP * GDP per capita as an approach that captures power through net resources rather than gross resources. We calculated Gross domestic product (GDP) and GDP per capita from the World Bank [44], average values 2007-2016. We summarized protected extent (in percentage) within Crowards’ groups employing box plots. We modeled the marine protection extent along size and military power gradients employing a Local Regression (LOESS) method. This non-parametric approach identifies patterns and fits a smoothed curve neither assuming any global function nor estimating a statistical significance of relationships (e.g., via a determination coefficient) [46].

In order to explore if protection is biased towards peripheral territories, i.e., geographically detached from the central territory, we categorized each EEZ into three categories: “Central territories” –which are the EEZs adjacent to the mainland of the sovereign country, “Undisputed peripheral territories” –which are the EEZs of those peripheral territories that are indisputably bounded to a sovereign country, and “Disputed peripheral territories” –which are the EEZs of those peripheral territories that overlap claims from two or more countries. Peripheral territories include first-level subnational units constituent of a sovereign country as well as other territories under different legal figures. Results are presented for each of the encompassed countries (i.e., a central territory and its associated peripheral territories are considered as a single unit), and individually for each of the EEZs. We performed a generalized linear model (GLM) to analyze the differences in the protected extent between the three EEZ categories. We used a quasi-Poisson error structure due to over-dispersion in the response variable [47] and performed a Tukey test in cases of significant differences in the model. A p-value of 0.001 was used as a threshold for statistical significance. Disputed peripheral territories were assigned to the *de facto* ruling country according to the Flanders Marine Institute database. Summarized national data are available in Appendix A, while summarized data of peripheral territories are available in Appendix B.

## 3. Results

The distribution of MPAs widely varies among countries. The micro and small countries of Cook Islands and Palau in the Pacific Ocean achieve the highest conservation values, with 100% and 80% of protection of their EEZs, respectively. Following them, Germany, the United Kingdom, the United States, Chile, France, and Australia exceeds 35% of the protected extent. It is worth noting that the United States, France, and Australia are the three countries with the largest EEZ area (Appendix A). Meanwhile, out of the 155 considered countries, 79 do not reach the 1% of protected extent, among them Brunei, Iceland, or the Federal States of Micronesia, with very different geographical and socio-economic characteristics. Notably, only 28 countries reach the protected 10% set by Target 11 of the Aichi agreement [5].

Conservation in the ocean tends to increase with Crowards’ group size (Figure 2a), and according to LOESS models, with land and ocean (EEZ) areas (Figure 2b-c), as well as with the three measurements of military power (Figure 2d-f). In particular, out of the 42 large to very large countries, nine exceed the 20% of the protected extent of their EEZs, while out of the 53 micro-to small-size countries, only two exceed this value (i.e., the above-mentioned Cook Islands and Palau). LOESS models show a notable increase in protection with increases in ocean area and GDP * GDP per capita, though the positive relationships can be observed especially towards the upper end of the five continuous gradients (Figure 2b-f).

**Figure 2.**
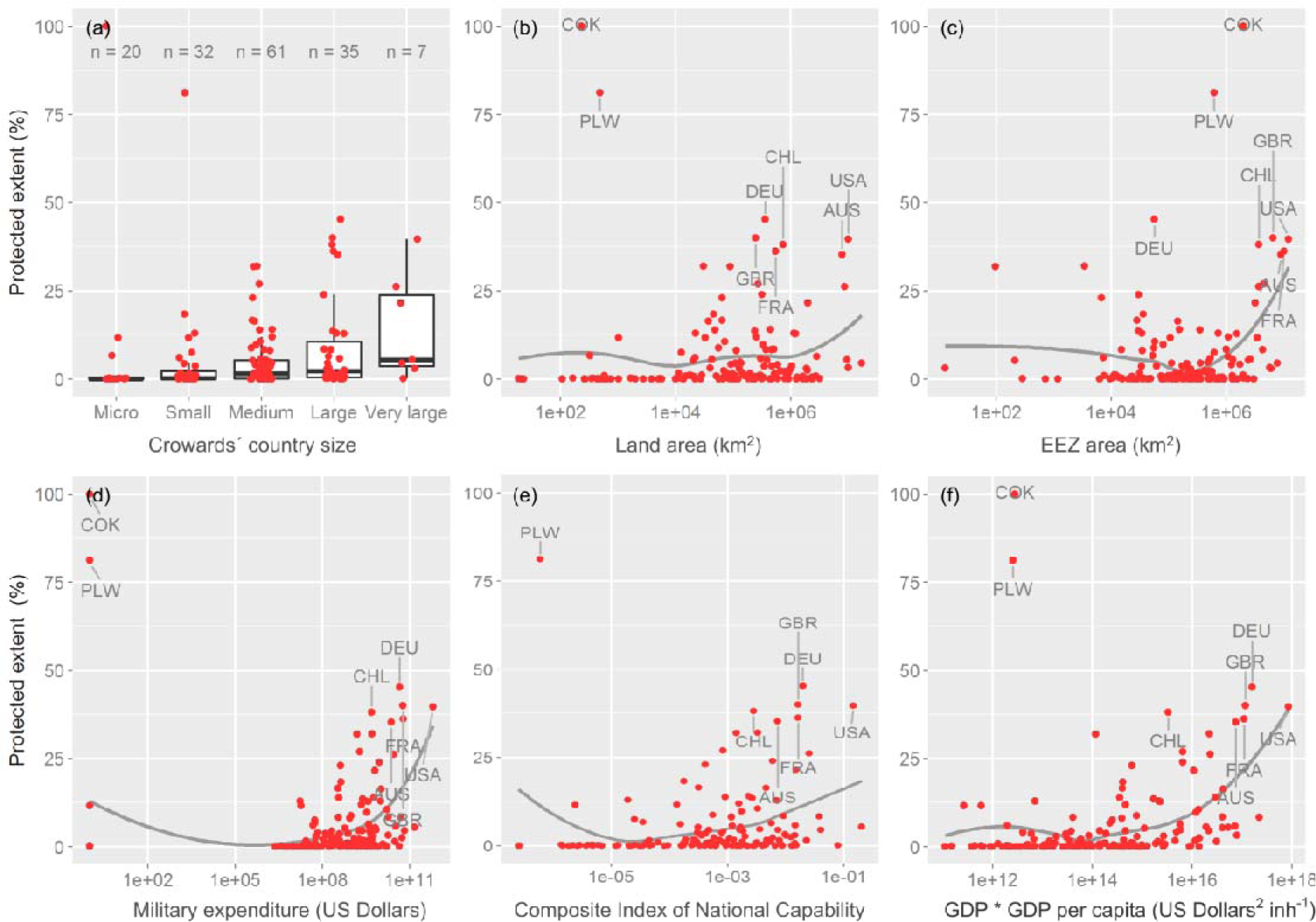
Marine protected extent (a) in groups of countries by size according to Crowards [35, 36], and (b-f) along gradients of size and military power. Peripheral territories are assigned to a Central territory. In (a) the extremes of the boxes represent the 25th and 75th quantiles; the horizontal line, the 50th quantile; whiskers, the hinge ± 1.5 * interquantile range; and red points the EEZ individual values. Countries with a protected extent > 35% are labeled in plots. Acronyms: Australia (AUS), Chile (Chile), Cook Islands (COK), Germany (DEU), France (FRA), United Kingdom (GBR), Palau (PLW), and United States of America (USA).

In light of these results, we wonder if the conservation extent of countries is influenced by the possession and effective management of peripheral territories. At a first cartographic glance, the peripheral territories host extensive MPAs (Figure 1b), both of disputed or undisputed sovereignty. Together with Cook Islands and Palau, they are, therefore, the EEZs of oceanic islands in the five oceans the ones that reach a higher protected extent (e.g., Easter Island, – Brazilian– Trindade and Martin Vaz, –French– Glorioso Islands) (Figure 1c). In quantitative terms, the analyzed 154 central territories have the statistically lowest protected extent value (mean = 6.4% ± 13.5, p-values < 0.001), in comparison to the other two types of analyzed territories, i.e., the disputed and undisputed peripheral territories, with similar mean and dispersion values (28.6% ± 40.4 and 20.8% ± 38.1, respectively; p-value = 0.493) (Figure 3a). Out of the 26 territories that overpass a 50% of protection, 18 are undisputed peripheral territories (e.g., Pitcairn Island, Guadeloupe), six are disputed peripheral territories (e.g., British Indian Ocean Territory), and only two central, or in this case, sovereign countries (i.e., the above-mentioned Cook Islands and Palau) (Appendixes A and B).

**Figure 3.**
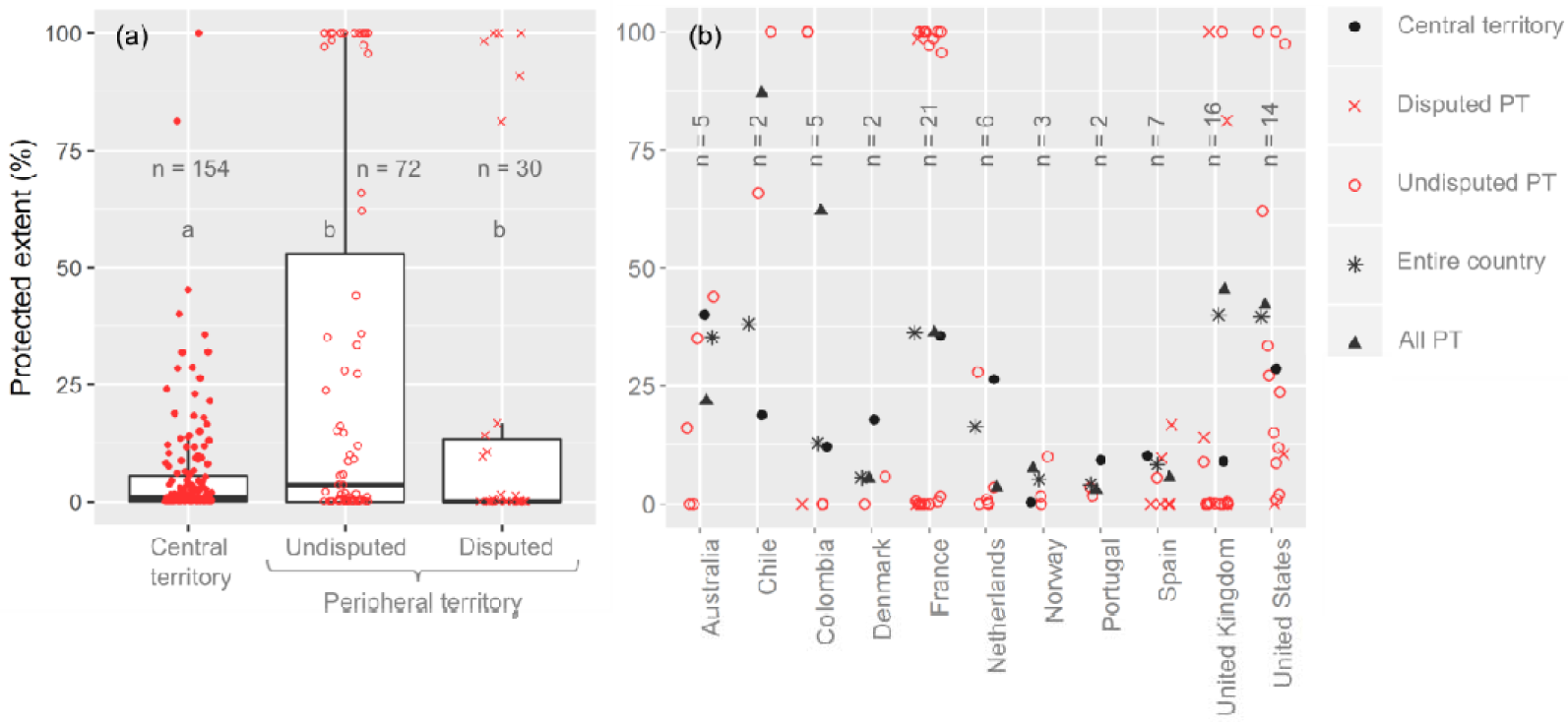
(a) Protection extent according to the category of the territory in the 256 EEZ considered. (b) Protection extent patterns for individual countries with two or more peripheral territories (with and without sovereignty disputes). The “entire country” category represents the protection extent of the sum of the areas of the central and peripheral territories. “All PT (peripheral territories)” type represents the average of the protection extent of undisputed and disputed territories. In (a) the extremes of the boxes represent the 25th and 75th quantiles; the horizontal line, the 50th quantile; whiskers, the hinge ± 1.5 * interquantile range; red points, the EEZ individual values; and letters, the significant differences between categories according to Tukey test (p < 0.001).

When these peripheral territories are linked graphically with their central territory, contrasting patterns arise between countries (Figure 3b). There are countries for which the conservation of the peripheral territories altogether is –in relative terms– higher than that of the central territory, such as the United States, the United Kingdom, Chile, Colombia, and to a lesser extent, Norway (Figure 3b). There are countries with the opposite pattern, i.e., the central territory has a higher level of conservation than its associated peripheral territories, such as Australia, Denmark, the Netherlands, and to a lesser extent Portugal and Spain. France is a unique case, as it is the only country in which the efforts of conservation of their coastal and marine territories are similarly distributed between the central territory and the set of peripheral territories. However, among these French peripheral territories (n = 21), there is a notable divergence, with approximately half reaching a high protected extent (> 95%; e.g., Martinique), and with the other half having a very low protected extent (< 1.6%; e.g., Wallis and Futuna).

## 4. Discussion

In this work, we found that it is some of the most powerful countries in the world that protect the ocean with greater intensity in terms of relative extent according to the Crowards’ classification [35, 36] and to LOESS models (Figure 2). Countries like the United States, the United Kingdom, France, Australia, or Chile, achieving a protection extent > 35% of their Exclusive Economic Zones (EEZs), are at the forefront of the conservation of the ocean through –mostly– very large marine protected areas (> 100,000 km^2^). The levels of protection reached by these countries surpass the targets set in Aichi [5] and forthcoming biodiversity agreements which suggest a rise from the current 10% to 30% by 2030 [48]. These results acquire particular relevance considering that these countries exert control over an extensive ocean surface (Figure 2c) and that they have the human, technological, legal, and financial resources to carry out concrete conservation actions [10]. Complementarily, medium to small countries or with little power might seem to be relegated in the process of conserving their oceanic biodiversity and resources. It is worth noting that the results summarized in the boxplot and LOESS models of Figure 2 emerge from 155 sovereign countries, therefore, the generally positive relationship between national power and ocean conservation would exceed the aforementioned particular cases.

With the global highest values of protection, the semi-independent Cook Islands and the sovereign Palau could be considered exceptions to the above-described general patterns, given their smallness in terms of land area, GDP, population, and military power. However, according to Chan [19], these and other countries in the Pacific and Indian Oceans should be named “Large Ocean States” instead of “Small Island Developing States” due to their extensive EEZs (Figure 2c). This author maintains that Large Ocean States are experiencing radical adjustments in the way of demonstrating their sovereignty over large tracts of the ocean, recognizing that size matters in geopolitics as a measure of power. So, although tentative, that the most powerful countries are the ones that protect the most, could be explained by the comparatively low *per capita* costs of public goods and services [49], their extensive human and economic resources [10], the pursue or competition for a moral leadership in the protection of marine environments [13], and as a way of effective presence, occupation, and territorial management and planning of the vast extensions under their jurisdiction [50] (concepts that we develop below), taking into consideration that the ocean is fundamentally under public rather than private tenure and control [51].

There are, however, several cases that deviate from Figure 2 models. At the high end of the national power gradients, countries such as Canada, China, India, Indonesia, or Russia, accounting together for 16.5% of the analyzed global EEZ area, protect a modest fraction of their coastal and marine territories (< 5.5%). This divergence between powerful countries had also been described by Shugart-Schmidt et al. [16], who challenged the entrenched idea that the countries with the greatest economic size or development are those that protect more and more efficiently their natural assets in the ocean [15, 52]. According to our results, a possible reason for this last divergence would be the lack of peripheral territories, since we found that these are foci of conservation actions (Figure 3 and Appendix B) [8, 16, 25]. In fact, out of the eleven countries that have two or more peripheral territories, six have a bias of the MPAs towards these areas (including the marginal case of France^5^). It is worth noting that in this paper we used a rather conservative concept of what might be called a “periphery”, as many geographically isolated territories were considered as part of the metropolises, including archipelagos at a distance less than 400 NM from the coast of a central territory (e.g., New Zealand Kermadec, Japanese Ryūkyū), as well as continental areas (e.g., Argentinean and Chilean Patagonia, Russian Siberia).

A first hypothesis regarding this geographical bias is that the designation of these peripheral territories as MPAs would imply relatively few political costs due to their remoteness with regards to the metropolis, null or low population (and therefore, voters) and infrastructure, low traffic, and poor economic importance or development [9, 12, 16]. As Jones and De Santo [54] posed, the main economic actors affected by the unilateral action of MPAs designations in peripheral territories eventually are the fishing or mining companies from foreign countries. In this way, it could be argued that countries with currently “idle” marine territories would follow an opportunistic rather than systematic approach to conservation in pursuit of compliance with international biodiversity standards or agreements such as those of Aichi [55, 56].

But this opportunistic approach to conservation is probably jointly acting with a group of geopolitical factors that are less explored and recognized by researchers as well as other social or professional communities [14]. Therefore, a second group of hypotheses is related to the geopolitical advantages that confer to a piece of ocean to be designated as a protected area. First and fundamentally, the creation of MAPs could be understood as a politically correct argument given by the largest and more powerful countries to justify and warrant their effective presence on any territory, which can be ultimately tagged as “global commons” [30]. This environmental and conservationist argument advocated by the administration of the central territory would change the opinion of the metropolitan citizens regarding the occupation of the ocean by essentially invoking their post-materialist values [57, 58]. At the same time, these national conservation policies are reinforced by receiving support from international NGOs in particular, and the international community (e.g., other governments) in general [10]. It is necessary to note that this last hypothesis would support both the conservation biases towards peripheral territories and the greater protection with the increase in the national power of countries.

Second, the conservationist argument also reconfigures the relationships between the central territory and their peripheral citizens (if land is inhabited), by potentially excluding local population from management decisions, denying their circulation through land and ocean, reducing the possibilities of populating the territory or even making them renounce their sovereign rights, that is, halting decolonizing and installing a contemporary political form of colonialism [21, 22].

Third, MPAs could be a mean to assure the provision of future goods and services for the central territory and its metropolitan population, even though currently many remote units have reduced importance providing assets [52]. As Ramutsindela et al. [27] affirmed, under the globally dominant economic/political logic, nature is often enclosed within legal boundaries in order to reserve areas of natural resources for future needs. This could be conceived as a “blue grabbing”, an extrapolation towards oceans [59] of the land/green grabbing concepts coined for continents [60].

Fourth and last, with the metropolitan or international supports assured, MPAs become an instrument that enforces the principle of territorial adjacency [20, 26]. This means to occupy the geographical space and project the influence of the metropolis over multiple geographical spaces. It is not a hidden fact that many peripheral territories house military facilities – including proving grounds– of confirmed strategic importance [29, 51]. For this reason, a few authors [e.g., 21, 25] refer that MPAs would be subject to “higher” interests and, consequently, effective conservation of biodiversity and ecosystem processes would not be secured in time. De Santo [25], in particular, highlights that in Marine Natural Monuments of the United States, presidential proclamations guarantee the military access when and where necessary. Regarding the expansion of state influence over global the ocean, Feral [20] details how groups of countries are increasingly agreeing to create numerous MPAs beyond national jurisdictions. As examples, within the framework of “Convention for the Protection of the Marine Environment of the North-East Atlantic”, 16 European contracting parties agreed to create MPAs in the Atlantic Ocean; while within the framework of the “Convention for the Conservation of Antarctic Marine Living Resources”, 26 members agreed to create a few MPAs in Antarctica while several new areas are being discussed [61]. Like in terms of global maritime security [30], these decisions from a reduced body of actors set important ocean policy precedents for the management or appropriation of resources over extra-jurisdictional territory.^6^

Probably, the above-mentioned hypotheses are not suitable for all peripheral territories and all times, as the encompassed EEZs vary widely in terms of their legal status, from semi-independent countries (e.g., Curaçao, Greenland), to subnational units (e.g., Hawaii, Réunion), to more uncertain cases (e.g., Clipperton, Jan Mayen), as well as from unpopulated (e.g., Macquarie Island) to densely populated (e.g., Puerto Rico) territories (Appendix B). However, certain paradigmatic cases reflect how, under conservationist and scientific aspirations, conservation, militarization, and geopolitics collide. The Chagos Islands, in the Indian Ocean, is the last oversea territory of the United Kingdom in Africa. In 1965, this country divided the colony of Mauritius for the probable purpose of creating a new administrative unit that would preserve the strategic position [21, 25]. This division openly challenged various resolutions of the United Nations General Assembly on Decolonization [22]. After the small indigenous population was expelled to facilitate the development and functioning of a major military base of the United States, the archipelago was designated as the British Indian Ocean Territory MPA in 2010. The designation of the MPA has drawn widespread criticism among human rights NGOs, the community of islanders in exile, and from the Mauritius administration [23, 24], as well as widespread support from environmental NGOs like Greenpeace or the Pew Environment Group. According to De Santo et al. [23], this “top-down” procedure would ring colonialism, while Sand [22] and many others maintain that the motivation for the designation of the MPA was to impede future resettlement of native inhabitants and their descendants in the territory and therefore to assure the strategic position.^7^

Finally, the role of geopolitical motivations conditioning or triggering the creation of MPAs is certainly open to discussion, as it cannot be easily isolated from the effects of other interacting forces [55, 56]. However, a key to elucidating the importance of geopolitics would be provided by the inspection of land conservation patterns. First, if biocentric motivations would guide national conservation agendas, and conservation differences were given exclusively by cultural or economic factors [57, 63], countries should maintain a similar position in the land and ocean conservation rankings. Nevertheless, as less powerful countries achieve nowadays the highest values of land protection [64], the contrasting land/ocean patterns would support the active role of geopolitical factors in ocean conservation. Second, geopolitical motivations seem to condition the early stages of conservation (in which ocean conservation probably is [56]). From the end of the 19th century until the middle of the 20th century, many powerful countries promoted the creation of very large land protected areas close to international boundaries, in depopulated (or populated by indigenous peoples) territories, with low production value, and under public tenure [65-67]. Beyond their generally great scenic value [56], the initial motivations for the conservation of these peripheral spaces were the effective occupation of the land by the central governments, (re)establishing international boundaries, and granting to a force –like park rangers– control and surveillance duties [18, 27, 65, 68].

## 5. Conclusions

Even with marked differences, the most powerful states are actively expanding their networks of protected areas. This advance in conservation is commonly associated with post-materialist ideals widely publicized and accepted by society, such as the conservation of biological and biophysical diversities [28]. However, multiple studies find that MPAs are biased towards isolated and sparsely populated territories of low economic importance [8, 9, 12]. In this study, we corroborate this geographic bias by identifying that many countries have declared much of their peripheries as MPAs, allowing them to comply with the Aichi Biodiversity agreement [5]. Nonetheless, we consider incomplete to understand the opportunism as the single or predominant motivation for this geographical bias. Based on previous scientific essays [20, 21, 25-27, 54], we provide data that allow assigning geopolitical factors as co-responsible for the observed phenomenon. This fact would not be new in the history of conservation, since the most powerful nations openly proclaim geopolitical their visions over the oceans, by maintaining strategic spatial positions that warrant their economic, political, cultural, and ideological predominance on the international arena as well as their future food security [30].

In this way, we understand that the compliance of the ongoing ultimate objective of MPAs (i.e., the long term conservation and sustainable management of marine assets) would be subordinated to veiled and broadest national interests (domestic and global). Therefore, political, economic and environmental security changes could lead to reductions of conservation funding, as well as a downgrading, downsizing, and degazettement of MPAs, as happened with their terrestrial counterparts [56, 69]. Finally, we would like to acknowledge that this in no way means that some countries do not specifically seek to represent the diversity of local biota or biophysical conditions through continuous expansion of their MPA network and continuous reevaluations of management plans [9, 28]. However, the geopolitical dimension of ocean conservation matters for the highest national decision-making levels and therefore should explicitly be acknowledged by researchers and practitioners, considering the strengths and weaknesses that this might imply in the preservation of biodiversity, the regulation of climate, and the sustainable provision of ecosystem services.

## Acknowledgements

We would like to thank J.I. Whitworth Hulse, T. Milani, S. Aguiar, E.G. Jobbágy for their ideas and collaboration in different stages of the study. This work was funded by a grant from the Argentinean National Scientific and Technical Research Council - CONICET (Proyectos de Unidades Ejecutoras 22920160100037CO).

## Footnotes

1 Höhn [31] points out that the major figures of American geopolitics defined that “The objective of geopolitics is to analyze security problems in terms of geographic factors and to make policy recommendations based on this analysis. Geopolitics is a technique to reckon with geographic position and physical power. In other words, geopolitical analysis is focused on power politics in a geographical context”.

2 Originally 280, but we separated the EEZ corresponding to Minami-Tori-shima from the EEZ of the central territory of Japan.

3 The entire EEZ of Ukraine is considered a disputed territory in Flanders Marine Institute database, with Ukraine as the de facto ruling country. Accordingly, the database is composed of 155 sovereign countries and 154 central territories.

4 As in 2018 the Chinese central administration requested the discontinuance of public sharing of their geographical data [38], we used –exclusively for this country– the protected extent value from the national profile available at www.protectedplanet.net.

5 This parity could change with the designation of new very large marine protected areas in its peripheral territories [53].

6 From the developed ideas, it could be argued that the peripheral territories under dispute between countries might have a larger protected extent than those that are not. However, in this work, we find high variability among cases and did not find statistically significant differences between both categories of periphery. Possible causes of this similarity could be the marginality of many disputed EEZs in terms of size (ten of the 31 cases do not exceed 1,000 km^2^) and in terms of administrative or territorial organization (ten have an unknown status and nine do not have a name assigned at the database of the Flanders Marine Institute) (Appendix B).

7 In February 2019, the United Nations International Court of Justice declared that the United Kingdom is under the obligation to end its administration of the Chagos Islands, as the territory had been illegitimately separated from Mauritius [62].

## References

[1] M. Coll, S. Libralato, S. Tudela, I. Palomera, F. Pranovi, Ecosystem overfishing in the ocean, PLoS ONE 3(12) (2008) e3881.

[2] M. Cole, P. Lindeque, C. Halsband, T.S. Galloway, Microplastics as contaminants in the marine environment: A review, Mar. Pollut. Bull. 62(12) (2011) 2588–2597.

[3] L.A. Barrie, D. Gregor, B. Hargrave, R. Lake, D. Muir, R. Shearer, B. Tracey, T. Bidleman, Arctic contaminants: Sources, occurrence and pathways, Sci. Total Environ. 122(1-2) (1992) 1–74.

[4] O. Hoegh-Guldberg, Climate change, coral bleaching and the future of the world’s coral reefs, Mar. Freshwater Res. 50(8) (1999) 839–866.

[5] SCBD, COP-10 Decision X/2. Secretariat of the convention on biological diversity, Nagoya, Japan. https://www.cbd.int/doc/decisions/cop-10/cop-10-dec-02-en.pdf, 2010.

[6] UNEP-WCMC, IUCN, NGS, Protected Planet Report 2018, Cambridge, UK; Gland, Switzerland; and Washington, D.C., USA. 2018.

[7] L. Boonzaier, D. Pauly, Marine protection targets: an updated assessment of global progress, Oryx 50(1) (2015) 27–35.

[8] L.J. Wood, L. Fish, J. Laughren, D. Pauly, Assessing progress towards global marine protection targets: Shortfalls in information and action, Oryx 42(3) (2008) 340–351.

[9] R. Devillers, R.L. Pressey, A. Grech, J.N. Kittinger, G.J. Edgar, T. Ward, R. Watson, Reinventing residual reserves in the sea: Are we favouring ease of establishment over need for protection?, Aquat. Conserv. 25(4) (2015) 480–504.

[10] J. Alger, P. Dauvergne, The politics of Pacific Ocean conservation: Lessons from the Pitcairn Islands Marine reserve, Pac. Aff. 90(1) (2017) 29–50.

[11] R.L. Singleton, C.M. Roberts, The contribution of very large marine protected areas to marine conservation: Giant leaps or smoke and mirrors?, Mar. Pollut. Bull. 87(1) (2014) 7–10.

[12] C. Kuempel, K. Jones, J. Watson, H. Possingham, Quantifying biases in marine-protected-area placement relative to abatable threats, Conserv. Biol. 33(6) (2019) 1350–1359.

[13] J. Alger, D. Peter, The global norm of large marine protected areas: Explaining variable adoption and implementation, Environ Policy Gov 27(4) (2017) 298–310.

[14] A.J. Caveen, T.S. Gray, S.M. Stead, N.V.C. Polunin, MPA policy: What lies behind the science?, Mar. Policy 37 (2013) 3–10.

[15] S. Marinesque, D.M. Kaplan, L.D. Rodwell, Global implementation of marine protected areas: Is the developing world being left behind?, Mar. Policy 36(3) (2012) 727–737.

[16] K.L.P. Shugart-Schmidt, E.P. Pike, R.A. Moffitt, V.R. Saccomanno, S.A. Magier, L.E. Morgan, SeaStates G20 2014: How much of the seas are G20 nations really protecting?, Ocean Coast. Manage. 115 (2015) 25–30.

[17] M. Fouqueray, E. Papyrakis, An empirical analysis of the cross-national determinants of marine protected areas, Mar. Policy 99 (2019) 87–93.

[18] T. Hodgetts, D. Burnham, A. Dickman, E.A. Macdonald, D.W. Macdonald, Conservation geopolitics, Conserv. Biol. 33(2) (2019) 250–259.

[19] N. Chan, “Large Ocean States”: Sovereignty, small Islands, and marine protected areas in global oceans governance, Global Gov. 24(4) (2018) 537–555.

[20] F. Feral, L’extension récente de la taille des aires marines protégées : une progression des surfaces inversement proportionnelle à leur normativité, VertigO - la revue électronique en sciences de l’environnement 9 (2011) 10998.

[21] P. Harris, Environmental protection as international security: Conserving the Pentagon’s island bases in the Asia–Pacific, Int J 69(3) (2014) 377–393.

[22] P.H. Sand, Fortress conservation trumps human rights?: The “Marine Protected Area” in the Chagos Archipelago, J. Environ. Dev. 21(1) (2012) 36–39.

[23] E.M. De Santo, P.J.S. Jones, A.M.M. Miller, Fortress conservation at sea: A commentary on the Chagos marine protected area, Mar. Policy 35(2) (2011) 258–260.

[24] P. Harris, Why law and politics matter for marine conservation - The case of the chagos marine protected area, Environ. Policy Law 45(5) (2015) 204–207.

[25] E.M. De Santo, Militarized marine protected areas in overseas territories: Conserving biodiversity, geopolitical positioning, and securing resources in the 21st century, Ocean Coast. Manage. 184 (2020) 105006.

[26] P. Leenhardt, B. Cazalet, B. Salvat, J. Claudet, F. Feral, The rise of large-scale marine protected areas: Conservation or geopolitics?, Ocean Coast. Manage. 85 (2013) 112–118.

[27] M. Ramutsindela, S. Guyot, S. Boillat, F. Giraut, P. Bottazzi, The geopolitics of protected areas, Geopolitics 25(1) (2019) 240–266.

[28] B.L. European Commission, and Agence française de développement, BEST Initiative. Projects 2011 - 2017, 28. https://ec.europa.eu/environment/nature/biodiversity/best/, 2017.

[29] N.J. Spykman, Geography and Foreign Policy, I, Am. Polit. Sc. Rev. 32(1) (1938) 28–50.

[30] B. Germond, The geopolitical dimension of maritime security, Mar. Policy 54 (2015) 137–142.

[31] K.H. Höhn, Geopolitics and the Measurement of National Power, Fachbereich Sozialwissenschaften, Universität Hamburg, Hamburg, Germany, 2011, p. 315.

[32] N. Prasad, Small but smart: small states in the global system, in: A.F. Copper, T.M. Shaw (Eds.), The Diplomacies of Small States. Between Vulnerability and Resilience, Palgrave Macmillan 2009, pp. 41–64.

[33] D. Immerwahr, How the US has hidden its empire, Farrar, Straus and Giroux, New York, US, 2019.

[34] J.F. Bradford, The maritime strategy of the United States: Implications for Indo-Pacific sea lanes, Contemp SE Asia 33(2) (2011) 183–208.

[35] T. Crowards, Defining the category of ‘small’ states, J. Int. Dev. 14(2) (2002) 143–179.

[36] T. Crowards, The comparative size of countries within Europe, Small States seminar, Manchester Metropolitan University, 2002, p. 11.

[37] Flanders Marine Institute, Maritime Boundaries Geodatabase: Maritime Boundaries and Exclusive Economic Zones (200NM), version 10, 2018. http://www.marineregions.org/. https://doi.org/10.14284/312.

[38] H.C. Bingham, D. Juffe Bignoli, E. Lewis, B. MacSharry, N.D. Burgess, P. Visconti, M. Deguignet, M. Misrachi, M. Walpole, J.L. Stewart, T.M. Brooks, N. Kingston, Sixty years of tracking conservation progress using the World Database on Protected Areas, Nat. Ecol. Evol. 3 (2019) 737–743.

[39] UNEP-WCMC and IUCN, Protected Planet: The World Database on Protected Areas (WDPA) March Release 2019 (web download version), Cambridge, UK. http://www.wdpa.org/, 2019.

[40] IUCN, Guidelines for Protected Area Management Categories, CNPPA with the assistance of WCMC, Gland, Switzerland and Cambridge, UK, 1994.

[41] M. Deguignet, A. Arnell, D. Juffe-Bignoli, Y. Shi, H. Bingham, B. MacSharry, N. Kingston, Measuring the extent of overlaps in protected area designations, PLOS ONE 12(11) (2017) e0188681.

[42] J.D. Singer, Reconstructing the correlates of war dataset on material capabilities of states, 1816–1985, Int. Interact. 14(2) (1988) 115–132.

[43] M. Beckley, The power of nations: Measuring what matters, Int. Security 43(2) (2018) 7–44.

[44] The World Bank, World Development Indicators, 2018. http://data.worldbank.org/.

[45] GlobalSecurity.org, World wide military expenditures, 2011.

[46] W.S. Cleveland, Robust locally weighted regression and smoothing scatterplots, J. Am. Stat. Assoc. 74(368) (1979) 829–836.

[47] A. Zuur, E.N. Ieno, N. Walker, A.A. Saveliev, G.M. Smith, Mixed effects models and extensions in ecology with R, Springer Science & Business Media 2009.

[48] SCBD, Zero draft of the post-2020 global biodiversity framework, Kunming, China. 14. 2020.

[49] A. Alesina, The Size of Countries: Does It Matter?, Journal of the European Economic Association 1(2-3) (2003) 301–316.

[50] L. Juda, Changing national approaches to ocean governance: The United States, Canada, and Australia, Ocean Dev. Int. Law 34(2) (2003) 161–187.

[51] J.-P. Lévy, A national ocean policy: An elusive quest, Mar. Policy 17(2) (1993) 75–80.

[52] H.E. Fox, C.S. Soltanoff, M.B. Mascia, K.M. Haisfield, A.V. Lombana, C.R. Pyke, L. Wood, Explaining global patterns and trends in marine protected area (MPA) development, Mar. Policy 36(5) (2012) 1131–1138.

[53] The Pew Charitable Trusts, French Polynesia’s Austral Islands Take Steps to Protect Their Waters. https://www.pewtrusts.org/en/research-and-analysis/articles/2016/04/05/french-polynesias-austral-islands-take-steps-to-protect-their-waters, 2016 (accessed November 2019).

[54] P.J.S. Jones, E.M. De Santo, Viewpoint – Is the race for remote, very large marine protected areas (VLMPAs) taking us down the wrong track?, Mar. Policy 73 (2016) 231–234.

[55] G. Baldi, M. Texeira, O.A. Martin, H.R. Grau, E.G. Jobbágy, Opportunities drive the global distribution of protected areas, PeerJ 5 (2017) e2989.

[56] J.E.M. Watson, N. Dudley, D.B. Segan, M. Hockings, The performance and potential of protected areas, Nature 515(7525) (2014) 67–73.

[57] Z. Baynham-Herd, T. Amano, W.J. Sutherland, P.F. Donald, Governance explains variation in national responses to the biodiversity crisis, Environ. Conserv. 45 (2018) 407–418.

[58] R. Inglehart, C. Welzel, Modernization, Cultural Change and Democracy, Cambridge University Press, New York, USA, 2005.

[59] N.J. Bennett, H. Govan, T. Satterfield, Ocean grabbing, Mar. Policy 57 (2015) 61–68.

[60] J. Fairhead, M. Leach, I. Scoones, Green Grabbing: a new appropriation of nature?, J. Peasant Stud. 39(2) (2012) 237–261.

[61] A.P. Capurro, Áreas marinas protegidas en Antártida: análisis de criterios para su designación, con énfasis en la región de la Península Antártica, Instituto Tecnológico de Buenos Aires, Buenos Aires, 2019, p. 93.

[62] The Guardian, UN court rejects UK’s claim of sovereignty over Chagos Islands. https://www.theguardian.com/world/2019/feb/25/un-court-rejects-uk-claim-to-sovereignty-over-chagos-islands, 2019 (accessed October 2019).

[63] P. Kashwan, Inequality, democracy, and the environment: A cross-national analysis, Ecol. Econ. 131 (2017) 139–151.

[64] G. Baldi, Nature protection across countries: Do size and power matter?, BioRxiv (2019).

[65] S. Marinaro, H.R. Grau, E. Aráoz, Extent and originality in the creation of national parks in relation to government and economical changes in Argentina, Ecología Austral 22(1) (2012) 1–10.

[66] C.L. Wirth, National Parks, in: A.B. Adams (Ed.), First World Conference on National Parks, National Park Service, Washington, USA, 1962, pp. 13–21.

[67] R.L. Pressey, Ad Hoc Reservations: Forward or Backward Steps in Developing Representative Reserve Systems?, Conserv Biol 8 (1994) 662–668.

[68] B. Sepúlveda, S. Guyot, A lo largo y a través de la frontera: áreas protegidas y gestión participativa en la Norpatagonia (Chile – Argentina), in: M.A. Nicoletti, P. Núñez, A. Núñez (Eds.), Araucanía-Norpatagonia: Discursos y representaciones de la materialidad, Editorial UNRN, IIDyPCa, Viedma, Argentina, 2016, pp. 243–273.

[69] M.B. Mascia, S. Pailler, Protected area downgrading, downsizing, and degazettement (PADDD) and its conservation implications, Conserv. Lett. 4(1) (2011) 9–20.

