## Appendix A for "Does national power trigger ocean conservation?"

| **Country** | **Crowards' country size** | **Land area**  (k km^2^) | **EEZ area,**  (k km^2^) | **Military expenditure**  (M US Dollars) | **Composite Indicator of National Material Capabilities** | **GDP * GDP per capita**  (M US Dollars^2^ inh^-1^) | **Protected extent of the EEZ** (%) |
| --- | --- | --- | --- | --- | --- | --- | --- |
| **Albania** | Sm | 28.34 | 12.15 | 183.6 | 0.0003 | 5.20E+07 | 1.06 |
| **Algeria** | La | 2308.86 | 131.14 | 7962.8 | 0.00404 | 8.70E+08 | 0.02 |
| **Angola** | Me | 1244.65 | 497.80 | 3849.1 | 0.00227 | 4.00E+08 | 0 |
| **Antigua and Barbuda** | Mi | 0.45 | 111.91 | 10 | 0 | 1.60E+07 | 0.17 |
| **Argentina*** | La | 2784.31 | 1069.36 | 3912.6 | 0.00568 | 5.80E+09 | 6.41 |
| **Australia*** | La | 7691.17 | 9000.32 | 23274.6 | 0.00728 | 7.50E+10 | 35.1 |
| **Azerbaijan** | Me | 86.25 | 80.52 | 2217.4 | 0.0014 | 3.60E+08 | 0.16 |
| **Bahamas** | Sm | 12.59 | 619.94 | 55 | 0.00002 | 2.50E+08 | 7.52 |
| **Bahrain** | Sm | 0.58 | 7.53 | 1051.9 | 0.00052 | 6.60E+08 | 1.01 |
| **Bangladesh** | La | 136.9 | 112.49 | 1868.7 | 0.00737 | 1.40E+08 | 4.47 |
| **Barbados** | Mi | 0.44 | 185.70 | 49.2 | 0.00002 | 7.40E+07 | 0.01 |
| **Belgium** | Me | 30.67 | 3.48 | 5165 | 0.00333 | 2.20E+10 | 31.9 |
| **Belize** | Sm | 22.3 | 34.43 | 17.2 | 0.00002 | 7.10E+06 | 13.0 |
| **Benin** | Me | 116.11 | 35.65 | 78.9 | 0.00041 | 6.60E+06 | 0 |
| **Bosnia and Herzegovina** | Me | 40.03 | 0.01 | 201.4 | 0.00038 | 8.50E+07 | 3.11 |
| **Brazil*** | VL | 8472.66 | 3690.32 | 29054.9 | 0.02511 | 2.30E+10 | 26.1 |
| **Brunei** | Sm | 5.72 | 43.33 | 412.6 | 0.00016 | 5.80E+08 | 0.03 |
| **Bulgaria** | Me | 112.76 | 34.69 | 878 | 0.00117 | 3.90E+08 | 8.01 |
| **Cabo Verde** | Sm | 3.88 | 804.69 | 9.4 | 0.00002 | 5.80E+06 | 0 |
| **Cambodia** | Me | 181.06 | 48.89 | 203.4 | 0.00157 | 1.40E+07 | 0.13 |
| **Cameroon** | Me | 464.32 | 15.21 | 360.1 | 0.00116 | 4.10E+07 | 8.25 |
| **Canada*** | La | 9945.53 | 5732.90 | 18473.6 | 0.00982 | 7.90E+10 | 3.26 |
| **Chile*** | La | 736.6 | 3668.54 | 4911.4 | 0.0028 | 3.30E+09 | 38.1 |
| **China*** | VL | 9374.57 | 1305.16 | 148000 | 0.20345 | 5.30E+10 | 5.41^[1]^ |
| **Colombia*** | La | 1135.12 | 731.82 | 10127.4 | 0.0068 | 2.10E+09 | 12.9 |
| **Comoros*** | Mi | 1.67 | 165.13 | 21.4 | 0.00003 | 4.50E+05 | 0.22 |
| **Congo** | Me | 344.89 | 33.95 | 388.2 | 0.00042 | 2.90E+07 | 3.55 |
| **Cook Islands** | Mi | 0.24 | 1976.46 | 0 | - | - | 100 |
| **Costa Rica** | Me | 51.14 | 584.22 | 293 | 0.00026 | 4.40E+08 | 0.83 |
| **Côte d'Ivoire** | Me | 320.68 | 172.52 | 425.1 | 0.00123 | 3.90E+07 | 0.09 |
| **Croatia** | Me | 55.08 | 55.30 | 972.9 | 0.00056 | 7.70E+08 | 8.72 |
| **Cuba** | Me | 109.93 | 353.20 | 2376.5 | 0.00162 | 4.60E+08 | 4.49 |
| **Cyprus** | Sm | 5.4 | 98.47 | 412.1 | 0.00017 | 6.70E+08 | 0.13 |
| **Dem. Rep. of the Congo** | La | 2325.24 | 13.43 | 281.1 | 0.00436 | 1.10E+07 | 0.48 |
| **Denmark*** | Me | 42.7 | 2619.39 | 4185.6 | 0.00119 | 1.90E+10 | 5.63 |
| **Djibouti*** | Sm | 21.85 | 7.25 | 48.7 | 0.00015 | 2.00E+06 | 0 |
| **Dominica** | Mi | 0.77 | 28.65 | 5.2 | 0 | 3.60E+06 | 0.01 |
| **Dominican Republic*** | Me | 48.44 | 351.76 | 378.2 | 0.00091 | 3.50E+08 | 13.5 |
| **Ecuador*** | Me | 255.01 | 1082.54 | 2233.1 | 0.00156 | 4.60E+08 | 11.7 |
| **Egypt*** | La | 1001.08 | 243.59 | 4768 | 0.0102 | 7.80E+08 | 2.8 |
| **El Salvador** | Me | 20.54 | 95.47 | 212.1 | 0.00043 | 9.10E+07 | 0.69 |
| **Equatorial Guinea** | Sm | 26.67 | 305.49 | 223.6 | 0.00005 | 3.00E+08 | 0.23 |
| **Eritrea*** | Sm | 122.54 | 78.63 | 445.8 | 0.00211 | 9.90E+05 | 0.04 |
| **Estonia** | Sm | 45.82 | 36.26 | 450.5 | 0.00018 | 4.00E+08 | 18.4 |
| **Federal States of Micronesia** | Mi | 0.63 | 3023.48 | 0 | 0.00001 | 9.20E+05 | 0.02 |
| **Fiji** | Sm | 18.93 | 1288.14 | 56.5 | 0.00006 | 1.80E+07 | 0.93 |
| **Finland** | Me | 333.06 | 81.07 | 3467.6 | 0.00173 | 1.20E+10 | 9.75 |
| **France*** | La | 547.84 | 10088.19 | 61724 | 0.01636 | 1.10E+11 | 36.2 |
| **Gabon** | Me | 259.97 | 202.65 | 227.1 | 0.00018 | 1.40E+08 | 0.69 |
| **Gambia** | Sm | 10.5 | 23.18 | 4 | 0.00009 | 1.60E+05 | 0.14 |
| **Georgia** | Me | 69.57 | 22.91 | 592.7 | 0.00046 | 5.30E+07 | 0.76 |
| **Germany** | La | 357.67 | 56.51 | 45222.2 | 0.01969 | 1.60E+11 | 45.2 |
| **Ghana** | Me | 238.67 | 228.51 | 188.8 | 0.0012 | 5.50E+07 | 0 |
| **Greece** | Me | 131.35 | 482.78 | 7104 | 0.0036 | 6.00E+09 | 4.69 |
| **Grenada** | Mi | 0.35 | 25.67 | 0 | 0 | 7.10E+06 | 0.08 |
| **Guatemala** | Me | 108.81 | 111.12 | 209.2 | 0.00083 | 1.70E+08 | 0.93 |
| **Guinea** | Me | 244.3 | 102.59 | 170 | 0.00057 | 5.10E+06 | 0.57 |
| **Guinea Bissau** | Sm | 32.83 | 107.30 | 18.6 | 0.00013 | 5.90E+05 | 11.6 |
| **Guyana*** | Sm | 211.21 | 139.25 | 35.7 | 0.00003 | 1.00E+07 | 0 |
| **Haiti*** | Sm | 26.89 | 117.75 | 6.4 | 0.00061 | 5.70E+06 | 1.25 |
| **Honduras** | Sm | 112.24 | 209.34 | 215.4 | 0.00048 | 3.80E+07 | 4.36 |
| **Iceland** | Me | 102.39 | 758.34 | 23.7 | 0.00003 | 8.00E+08 | 0 |
| **India*** | VL | 3151.45 | 2332.56 | 44648.4 | 0.07943 | 2.60E+09 | 0.12 |
| **Indonesia** | VL | 1879.83 | 6051.53 | 5682.1 | 0.01441 | 2.70E+09 | 2.97 |
| **Iran*** | La | 1622.51 | 215.58 | 11985.2 | 0.01448 | 2.80E+09 | 0.73 |
| **Iraq** | Me | 437.37 | 1.19 | 5027.9 | 0.00662 | 9.70E+08 | 0 |
| **Ireland** | Me | 69.45 | 425.35 | 1266.8 | 0.00058 | 1.40E+10 | 2.37 |
| **Israel** | Me | 21.9 | 24.64 | 15619.4 | 0.0041 | 8.90E+09 | 0.05 |
| **Italy** | La | 301.18 | 536.13 | 34069 | 0.01451 | 7.30E+10 | 5.79 |
| **Jamaica** | Sm | 11.03 | 257.78 | 119.7 | 0.00018 | 6.80E+07 | 0.68 |
| **Japan*** | La | 373.51 | 4286.43 | 49562.4 | 0.03737 | 2.20E+11 | 7.88 |
| **Jordan** | Me | 88.86 | 0.10 | 1547 | 0.00142 | 1.20E+08 | 31.8 |
| **Kazakhstan** | La | 2714.27 | 114.14 | 1780.4 | 0.00321 | 1.90E+09 | 0.9 |
| **Kenya*** | Me | 585.7 | 164.79 | 733.5 | 0.00172 | 5.90E+07 | 0.56 |
| **Kiribati** | Mi | 1.03 | 3455.26 | 0 | 0 | 2.70E+05 | 11.6 |
| **Kuwait** | Me | 17.47 | 11.19 | 2346.7 | 0.00179 | 2.50E+09 | 1.11 |
| **Latvia** | Me | 64.58 | 28.21 | 347.6 | 0.00032 | 4.00E+08 | 16.5 |
| **Lebanon** | Me | 10 | 20.19 | 1725.2 | 0.00094 | 3.50E+08 | 0 |
| **Liberia** | Sm | 95.3 | 252.89 | 10.3 | 0.00024 | 6.50E+05 | 0.07 |
| **Libya** | Me | 1623.76 | 364.70 | 722.7 | 0.00147 | 5.80E+08 | 0 |
| **Lithuania** | Me | 64.94 | 6.80 | 428.9 | 0.00041 | 6.20E+08 | 23 |
| **Madagascar*** | Me | 592.98 | 1243.22 | 72.5 | 0.00084 | 4.10E+06 | 4.06 |
| **Malaysia** | La | 327.88 | 513.04 | 4494.6 | 0.00461 | 2.80E+09 | 0.32 |
| **Maldives** | Mi | 0.11 | 929.34 | 92.6 | 0.00001 | 2.50E+07 | 0.05 |
| **Malta** | Mi | 0.33 | 52.92 | 55.4 | 0.00004 | 2.30E+08 | 6.66 |
| **Marshall Islands** | Sm | 0.16 | 2009.62 | 0 | 0 | 5.90E+05 | 0.28 |
| **Mauritania*** | Sm | 1036.39 | 173.73 | 131.5 | 0.0003 | 6.00E+06 | 3.72 |
| **Mauritius*** | Me | 2.01 | 1282.42 | 17.7 | 0.00005 | 1.00E+08 | 0 |
| **Mexico** | VL | 1957.85 | 3195.53 | 6373.1 | 0.01512 | 1.10E+10 | 21.5 |
| **Monaco** | Mi | 0.02 | 0.29 | 0 | 0 | 9.10E+08 | 0.14 |
| **Montenegro** | Sm | 13.73 | 6.37 | 70.4 | 0.00008 | 3.00E+07 | 0.01 |
| **Morocco*** | La | 591.74 | 280.60 | 3289 | 0.00388 | 2.90E+08 | 0.16 |
| **Mozambique** | Me | 788.45 | 567.88 | 116.3 | 0.00109 | 6.70E+06 | 2.19 |
| **Myanmar** | La | 663.16 | 499.28 | 1924.2 | 0.006 | 5.90E+07 | 2.28 |
| **Namibia** | Me | 822.71 | 563.51 | 393.2 | 0.00019 | 5.80E+07 | 1.71 |
| **Nauru** | Mi | 0.02 | 310.65 | 0 | 0 | 6.30E+05 | 0 |
| **Netherlands*** | Me | 37.1 | 144.90 | 10751.8 | 0.00461 | 4.20E+10 | 16.4 |
| **New Zealand*** | Me | 268.49 | 4420.15 | 1972.7 | 0.00083 | 6.50E+09 | 26.9 |
| **Nicaragua** | Me | 128.69 | 214.63 | 60.5 | 0.00036 | 1.90E+07 | 2.8 |
| **Nigeria** | La | 907.5 | 179.84 | 2104.2 | 0.0087 | 1.00E+09 | 0.02 |
| **Niue** | Sm | 0.26 | 319.09 | 0 | NA | #¡VALOR! | 0.01 |
| **North Korea** | La | 122.38 | 114.26 | 1000 | 0.01272 | 1.10E+07 | 0 |
| **Norway*** | Me | 382.07 | 2447.69 | 6619.1 | 0.00167 | 4.10E+10 | 5.28 |
| **Oman** | Me | 311.21 | 558.19 | 7667.1 | 0.0013 | 1.30E+09 | 0.12 |
| **Pakistan** | La | 872.94 | 224.90 | 7363.4 | 0.01419 | 2.70E+08 | 0.25 |
| **Palau** | Sm | 0.49 | 617.45 | 0 | 0 | 2.60E+06 | 81.2 |
| **Panama** | Me | 74.53 | 332.71 | 389.7 | 0.00027 | 4.20E+08 | 1.89 |
| **Papua New Guinea** | Me | 465.15 | 2409.92 | 73.2 | 0.00025 | 4.50E+07 | 0.18 |
| **Peru** | La | 1289.87 | 857.98 | 2213.8 | 0.00326 | 9.90E+08 | 0.47 |
| **Philippines** | La | 293.24 | 1978.55 | 2840.8 | 0.00546 | 6.10E+08 | 1.1 |
| **Poland** | La | 313.43 | 29.85 | 9245.9 | 0.00598 | 6.50E+09 | 24 |
| **Portugal*** | Me | 91.39 | 1728.04 | 4399.3 | 0.00184 | 4.90E+09 | 4.17 |
| **Qatar*** | Sm | 11.15 | 31.55 | 2788.3 | 0.00126 | 1.20E+10 | 1.72 |
| **Romania** | La | 236.38 | 29.55 | 2486.3 | 0.00266 | 1.70E+09 | 13.5 |
| **Russia** | VL | 16953.09 | 7686.38 | 68121 | 0.04021 | 2.10E+10 | 4.51 |
| **Saint Kitts and Nevis** | Mi | 0.27 | 9.53 | 0 | 0 | 1.20E+07 | 0.05 |
| **Saint Lucia** | Mi | 0.61 | 15.47 | 0 | 0 | 1.20E+07 | 0.28 |
| **Saint Vincent and the Grenadines** | Mi | 0.37 | 36.38 | 0 | 0 | 4.60E+06 | 0.23 |
| **Samoa** | Mi | 2.78 | 130.97 | 0 | 0 | 2.90E+06 | 0.01 |
| **São Tomé and Príncipe** | Mi | 1.04 | 131.43 | 2.4 | 0.00001 | 3.80E+05 | 0 |
| **Saudi Arabia** | La | 1921.73 | 224.78 | 56042.6 | 0.01247 | 1.40E+10 | 2.46 |
| **Senegal** | Me | 196.22 | 158.94 | 219.8 | 0.00073 | 1.40E+07 | 1.06 |
| **Seychelles** | Mi | 0.49 | 1347.25 | 13.7 | 0 | 1.60E+07 | 0.02 |
| **Sierra Leone** | Sm | 71.61 | 161.28 | 32 | 0.00032 | 1.90E+06 | 0.24 |
| **Singapore** | Me | 0.51 | 0.72 | 8733.5 | 0.00291 | 1.40E+10 | 0 |
| **Slovenia** | Me | 20.33 | 0.21 | 604.9 | 0.00031 | 1.10E+09 | 5.23 |
| **Solomon Islands** | Sm | 27.2 | 1611.84 | 44.8 | 0.00001 | 1.60E+06 | 0.06 |
| **Somalia** | Me | 471.82 | 785.21 | 51 | 0.00063 | 2.40E+06 | 0 |
| **South Africa*** | La | 1219.82 | 1546.75 | 3839.4 | 0.00723 | 2.30E+09 | 12.8 |
| **South Korea** | Me | 98.54 | 347.64 | 31440.7 | 0.0228 | 3.10E+10 | 3.48 |
| **Spain*** | La | 506.88 | 1007.62 | 18450.8 | 0.00922 | 4.00E+10 | 8.37 |
| **Sri Lanka** | Me | 66.29 | 535.85 | 1720.2 | 0.0021 | 2.20E+08 | 0.06 |
| **Sudan*** | Me | 1857.64 | 82.80 | 3140.1 | 0.00276 | 1.40E+08 | 4.38 |
| **Suriname** | Sm | 145.12 | 133.87 | 27.8 | 0.00004 | 3.60E+07 | 1.56 |
| **Sweden** | Me | 446.17 | 154.51 | 5982.5 | 0.0023 | 2.90E+10 | 14 |
| **Syria** | La | 185.94 | 10.27 | 1656.8 | 0.00454 | 8.30E+07 | 0.23 |
| **Tanzania** | Me | 941.51 | 242.60 | 342.7 | 0.00201 | 3.10E+07 | 2.67 |
| **Thailand** | La | 514.45 | 299.92 | 5222.4 | 0.00778 | 2.10E+09 | 1.47 |
| **Timor-Leste** | Sm | 15.08 | 74.31 | 32.3 | 0.00004 | 1.30E+06 | 0.78 |
| **Togo** | Sm | 56.86 | 15.52 | 64.2 | 0.00033 | 2.10E+06 | 0 |
| **Tonga** | Mi | 0.6 | 667.96 | 6.8 | 0 | 1.60E+06 | 0.02 |
| **Trinidad and Tobago*** | Sm | 5.12 | 76.87 | 171.7 | 0.00051 | 4.40E+08 | 0.02 |
| **Tunisia** | Me | 156.61 | 99.69 | 721.8 | 0.00095 | 1.80E+08 | 0.98 |
| **Turkey** | La | 780.08 | 261.93 | 17101.8 | 0.01518 | 9.40E+09 | 0.14 |
| **Turkmenistan** | Me | 470.85 | 61.17 | 787.5 | 0.00079 | 1.90E+08 | 3.65 |
| **Tuvalu** | Mi | 0.02 | 756.31 | 0 | 0 | 1.10E+05 | 0.01 |
| **Ukraine*^[2]^** | La | 599.09 | 135.90 | 4092.4 | 0.00927 | 4.40E+08 | 1.45 |
| **United Arab Emirates*** | Me | 71.09 | 57.97 | 16992 | 0.00332 | 1.30E+10 | 10.6 |
| **United Kingdom*** | La | 243.78 | 6552.21 | 58692.1 | 0.01645 | 1.20E+11 | 40.4 |
| **United States of America*** | VL | 9833.52 | 12160.59 | 646000 | 0.14573 | 8.50E+11 | 39.6 |
| **Uruguay** | Me | 177.34 | 143.08 | 809.2 | 0.00052 | 6.70E+08 | 0.61 |
| **Vanuatu** | Sm | 12.3 | 625.53 | 0 | 0.00001 | 2.20E+06 | 0.23 |
| **Venezuela*** | La | 912.68 | 475.47 | 4132.3 | 0.00472 | 4.60E+09 | 1.66 |
| **Vietnam** | La | 328.89 | 753.88 | 3243.9 | 0.00837 | 2.70E+08 | 0.45 |
| **Yemen** | Me | 453.07 | 529.39 | 1493.9 | 0.00168 | 4.50E+07 | 0 |

1. This value came from the UNEP-WCMC and IUCN country profile (www.protectedplanet.net). Otherwise, 0.32 from the geographical data of the World Database on Protected Areas, version March 2019.
2. The entire EEZ is considered a disputed territory in Flanders Marine Institute database. While not depicted in Appendix B, in analyses is treated as such.
