## Appendix B for "Does national power trigger ocean conservation?"

| **Country** | **Type** | **Territory** | **Status** | **EEZ area** (k km^2^) | **Protected extent of the EEZ** (%) |
| --- | --- | --- | --- | --- | --- |
| **Australia** | **CT** | **-** | **-** | **6881.49** | **40.10** |
|  | UT | Christmas Is. | External territory | 329.30 | 0.00 |
|  |  | Cocos Is. |  | 469.26 | 0.00 |
|  |  | Heard and McDonald Is. |  | 415.58 | 16.19 |
|  |  | Norfolk Is. |  | 431.17 | 43.92 |
|  |  | Macquarie Is. | Part of a subnational unit | 473.52 | 34.99 |
| **Brazil** | **CT** | **-** | **-** | **3216.50** | **14.95** |
|  | UT | Trindade and Martin Vaz | Part of a subnational unit | 473.82 | 100.00 |
| **Canada** | **CT** | **-** | **-** | **5708.32** | **3.28** |
| **Canada*/United States of America** | DT | Unnamed | Unknown | 24.58 | 0.00 |
| **Chile** | **CT** | **-** | **-** | **2485.71** | **18.88** |
|  | UT | Desventuradas Is. | Part of a subnational unit | 451.79 | 65.84 |
|  |  | Easter Is. | Special territory, part of a subnational unit | 731.04 | 100.00 |
| **China/Brunei/Indonesia/Malaysia/**  **Philippines/Taiwan/Vietnam** | DT | South China Sea† | Disputed territory | 340.12 | 0.00 |
| **Colombia** | **CT** | **-** | **-** | **721.28** | **12.12** |
|  | UT | Bajo Nuevo | Part of a subnational unit | 1.56 | 0.00 |
|  |  | Quitasueño |  | 2.85 | 100.00 |
|  |  | Serrana |  | 3.70 | 100.00 |
|  |  | Serranilla |  | 1.56 | 0.00 |
| **Colombia*/Dominican Republic/Venezuela** | DT | Unnamed | Unknown | 0.87 | 0.00 |
| **Denmark** | **CT** | **-** | **-** | **104.52** | **17.92** |
|  | UT | Faeroe | Autonomous territory | 262.56 | 0.01 |
|  |  | Greenland |  | 2252.31 | 5.73 |
| **Ecuador** | **CT** | **-** | **-** | **237.35** | **2.15** |
|  | UT | Galápagos | Province | 845.19 | 14.64 |
| **Eritrea** | **CT** | **-** | **-** | **78.38** | **0.00** |
| **Eritrea*/Djibouti** | DT | Doumeira Is. | Unknown | 0.25 | 0.00 |
| **France** | **CT** | **-** | **-** | **344.40** | **35.57** |
|  | UT | New Caledonia | Exceptional collectivity | 1179.24 | 95.56 |
|  |  | French Polynesian | Overseas collectivity | 4781.06 | 0.00 |
|  |  | Saint-Barthélemy |  | 4.19 | 98.42 |
|  |  | Saint-Martin |  | 1.10 | 97.11 |
|  |  | Saint-Pierre and Miquelon |  | 12.38 | 0.00 |
|  |  | Wallis and Futuna |  | 263.75 | 0.00 |
|  |  | French Guiana | Overseas department and region | 132.12 | 0.57 |
|  |  | Guadeloupe |  | 91.01 | 100.00 |
|  |  | Martinica |  | 47.96 | 100.00 |
|  |  | Réunion |  | 315.98 | 0.01 |
|  |  | Amsterdam & Saint Paul Is. | Overseas territory | 511.85 | 100.00 |
|  |  | Bassas da India |  | 121.04 | 0.00 |
|  |  | Crozet Is. |  | 573.93 | 100.00 |
|  |  | Europa Is. |  | 128.24 | 1.62 |
|  |  | Juan de Nova |  | 62.98 | 0.00 |
|  |  | Kerguelen |  | 549.39 | 100.00 |
|  |  | Clipperton | State property | 437.42 | 0.41 |
| **France*/Comoros** | DT | Mayotte | Overseas department and region | 66.99 | 98.38 |
| **France*/Madagascar** |  | Glorioso Is. | Overseas territory | 43.68 | 100.00 |
| **France*/Mauritius** |  | Tromelin Is. |  | 275.51 | 0.00 |
| **France*/Vanuatu** |  | Matthew and Hunter Is. | Part of New Caledonia | 187.63 | 100.00 |
| **Guyana** | **CT** | **-** | **-** | **135.63** | **0.00** |
| **Guyana*/Trinidad and Tobago/Venezuela** | DT | Unnamed | Unknown | 3.63 | 0.00 |
| **India** | **CT** | **-** | **-** | **1665.41** | **0.15** |
|  | UT | Andaman and Nicobar Is. | [Union territory](https://en.wikipedia.org/wiki/Union_territory) | 667.14 | 0.05 |
| **Japan** | CT | - | - | 3642.62 | 9.32 |
|  | UT | Minami-Tori-shima | Part of a subnational unit | 428.88 | 0.00 |
| **Kenya** | **CT** | **-** | **-** | **114.34** | **1.09** |
| **Kenya*/Somalia** | DT | Unnamed | Unknown | 50.45 | 0.00 |
| **Mauritania/Morocco/Polisario Front** | DT | Western Saharan† | Disputed territory | 284.73 | 1.31 |
| **Netherlands** | **CT** | **-** | **-** | **64.06** | **26.37** |
|  | UT | Aruba | Constituent country | 30.13 | 0.00 |
|  |  | Curaçao |  | 25.50 | 0.02 |
|  |  | Sint-Maarten |  | 0.47 | 3.59 |
|  |  | Bonaire | Special municipality | 13.05 | 0.20 |
|  |  | Saba |  | 9.52 | 27.94 |
|  |  | Sint-Eustatius |  | 2.18 | 0.98 |
| **New Zealand** | **CT** | **New Zealand** | **-** | **4098.26** | **28.44** |
|  | UT | Tokelau | Self-administering territory | 321.89 | 0.00 |
| **Norway** | **CT** | **-** | **-** | **926.63** | **0.40** |
|  | UT | Bouvet | Dependency | 439.91 | 0.00 |
|  |  | Svalbard | Exceptional unincorporated territory | 790.44 | 10.02 |
|  |  | Jan Mayen | Unincorporated territory | 290.71 | 1.63 |
| **Portugal** | **CT** | **-** | **-** | **315.29** | **9.30** |
|  | UT | Azores | Autonomous Region | 959.75 | 3.53 |
|  |  | Madeira |  | 453.01 | 1.81 |
| **Qatar** | **CT** | **-** | **-** | **31.42** | **1.36** |
| **Qatar*/Saudi Arabia/United Arab Emirates** | DT | Unnamed | Unknown | 0.13 | 90.91 |
| **Russia** | **CT** | **-** | **-** | **7686.38** | **4.51** |
| **Russia*/Japan** | DT | Kuril Is. | Part of a subnational unit | 213.31 | 0.50 |
| **South Africa** | **CT** | **-** | **-** | **1073.11** | **0.43** |
|  | UT | Prince Edward Is. | Part of a subnational unit | 473.64 | 35.79 |
| **South Korea** | **CT** | **-** | **-** | **347.64** | **1.49** |
|  | DT | Liancourt Rocks | Part of a subnational unit | 1.62 | 0.20 |
| **Spain** | **CT** | **-** | **-** | **560.96** | **10.27** |
|  | UT | Canary Is. | Autonomous community | 446.56 | 5.57 |
| **Spain*/Morocco** | DT | Ceuta | Autonomous city | 0.03 | 9.74 |
|  |  | Melilla |  | 0.03 | 0.00 |
|  |  | Alhucemas Is. | Places of sovereignty | 0.01 | 0.00 |
|  |  | Chafarinas Is. |  | 0.03 | 16.79 |
|  |  | Peñón de Vélez de la Gomera |  | 0.00 | 0.00 |
|  |  | Perejil Is. |  | 0.00 | 0.00 |
| **Sudan** | **CT** | **-** | **-** | **63.01** | **0.02** |
| **Sudan*/Egypt** | DT | Unnamed | Unknown | 19.78 | 0.00 |
| **United Kingdom** | **CT** | **-** | **-** | **729.12** | **9.03** |
|  | UT | Guernsey | Crown dependency | 6.51 | 0.26 |
|  |  | Jersey |  | 2.27 | 8.98 |
|  |  | Anguilla | Overseas territory | 90.43 | 0.04 |
|  |  | Ascension |  | 447.89 | 0.00 |
|  |  | Bermudas |  | 453.45 | 0.00 |
|  |  | British Virgin Is. |  | 81.79 | 0.00 |
|  |  | Cayman Is. |  | 118.65 | 0.08 |
|  |  | Montserrat |  | 7.21 | 0.00 |
|  |  | Pitcairn, Henderson, Ducie and Oeno Is. | | 844.11 | 100.00 |
|  |  | Saint Helena |  | 450.77 | 0.00 |
|  |  | Tristan Da Cunha |  | 757.59 | 0.52 |
|  |  | Turks and Caicos |  | 91.25 | 0.17 |
| **United Kingdom*/Argentina** | DT | Falkland Is. |  | 548.50 | 0.01 |
|  |  | South Georgia and South Sandwich | | 1281.44 | 81.14 |
| **United Kingdom*/Spain** |  | Gibraltar |  | 0.39 | 14.14 |
| **United Kingdom*/Mauritius** |  | British Indian Ocean Territory |  | 640.84 | 100.00 |
| **United Arab Emirates** | **CT** | **-** | **-** | **51.78** | **11.84** |
| **United Arab Emirates*/Iran** | DT | Unnamed | Unknown | 6.19 | 0.00 |
| **United States of America** | **CT** | **-** | **-** | **2452.57** | **28.63** |
|  | UT | Palmyra Atoll | Incorporated, unorganized territory | 355.19 | 15.14 |
|  |  | Alaska | State | 3664.69 | 33.44 |
|  |  | Hawaii |  | 2480.25 | 62.02 |
|  |  | Guam | Unincorporated, organized territory | 209.03 | 23.69 |
|  |  | Virgin Is. |  | 38.40 | 0.96 |
|  |  | Northern Mariana Is. | Unincorporated, organized territory (commonwealth) | 766.03 | 27.24 |
|  |  | Puerto Rico |  | 155.16 | 2.05 |
|  |  | American Samoa | Unincorporated, unorganized territory | 407.33 | 8.67 |
|  |  | Howland and Baker Is. |  | 436.84 | 11.88 |
|  |  | Jarvis Is. |  | 324.63 | 97.39 |
|  |  | Johnston Atoll |  | 443.93 | 100.00 |
|  |  | Wake Is. |  | 408.19 | 100.00 |
| **United States of America*/Haiti** | DT | Navassa Is. |  | 13.93 | 10.58 |
| **United States of America*/Dominican Republic** |  | Unnamed | Unknown | 18.35 | 0.00 |
| **Venezuela** | **CT** | **-** | **-** | **474.35** | **1.67** |
| **Venezuela*/Aruba/**  **Dominican Republic** | DT | Unnamed | Unknown | 1.12 | 0.00 |
